# Theta and gamma hippocampal–neocortical oscillations during the episodic-like memory test: impairment in epileptogenic rats

**DOI:** 10.1101/2021.12.09.471759

**Authors:** Anton Malkov, Ludmila Shevkova, Aleksandra Latyshkova, Valentina Kitchigina

## Abstract

Cortical oscillations in different frequency bands have been shown to be intimately involved in exploration of environment and cognition. Here, the local field potentials in the hippocampus, the medial prefrontal cortex (mPFC), and the medial entorhinal cortex (mEC) were recorded simultaneously in rats during the execution of the episodic-like memory task. The power of hippocampal theta (~4-10 Hz), slow gamma (~25-50 Hz), and fast gamma oscillations (~55-100 Hz) was analyzed in all structures examined. Particular attention was paid to the theta coherence between three mentioned structures. The modulation of the power of gamma rhythms by the phase of theta cycle during the execution of the episodic-like memory test by rats was also closely studied. Healthy rats and rats one month after kainate-induced status epilepticus (SE) were examined. Paroxysmal activity in the hippocampus (high amplitude interictal spikes), excessive excitability of animals, and the death of hippocampal and dentate granular cells in rats with kainate-evoked SE were observed, which indicated the development of seizure focus in the hippocampus (epileptogenesis). One month after SE, the rats exhibited a specific impairment of episodic memory for the *what-where-when* triad: unlike healthy rats, epileptogenic SE animals did not identify the objects during the test. This impairment was associated with the changes in the characteristics of theta and gamma rhythms and specific violation of theta coherence and theta/gamma coupling in these structures in comparison with the healthy animals. We believe that these disturbances in the cortical areas play a role in episodic memory dysfunction in kainate-treated animals. These findings can shed light on the mechanisms of cognitive deficit during epileptogenesis.

## Introduction

Oscillations represent the basic activity of the healthy brain, and the theta and gamma rhythms are crucial for cognition and memory (Buzsaki, 2006; Buzsáki and Draguhn, 2004; Lachaux, et al., 2005; Fries, 2005; Womelsdorf et al., 2006; Buzsaki and Watson, 2012). It was proposed that the theta rhythm is important for the reception and processing of new sensory signals and accomplishes their selection when comparing them with the information stored in the memory system (Vinogradova, 1995, 2001; Buzsáki, 2006; Colgin, 2013). It was also suggested that the theta rhythm (~4-10 Hz) is a critical mechanism for linking different event attributes into a single contextual representation, both in humans (Fell et al., 2001; Lega et al., 2012; Fell and Axmacher, 2011) and rodents (Inostroza et al., 2013b). The main function of the gamma rhythm is the performance of inter-regional communication and the selection of signals (Fries et al., 2007; Fries, 2009). Besides, gamma oscillations combine the activity of distributed neurons that process various properties of stimuli (for example, visual), and convert these properties into coherent perception (Gray et al., 1989). It is assumed that the mechanisms of organization of slow (~25-50 Hz) and fast (~55-100 Hz) gamma-oscillations are different since they are generated by different networks involving some particular classes of GABAergic interneurons (for review, see Colgin, 2016; Mably and Colgin, 2018).

External or internal events can lead to the synchronization of rhythms generated in the same or different brain regions and the formation of a more complex functional phenomenon known as phase coherence, or phase coupling (Fell et al., 2008; Cavanagh et al., 2009; Canolty and Knight, 2010). Standard phase coherence reveals the relative constancy of phase differences between two oscillations of the same frequency recorded in different regions, i.e., intra-frequency coherence (Rodriguez et al., 1999; Lachaux et al., 1999; Hurtado et al., 2004). It has been shown that phase coherence reflects various cognitive processes in humans (Canolty et al., 2006; Axmacher et al., 2010) and animals (Montgomery and Buzsáki, 2007; Montgomery et al., 2008; Tort et al., 2008, 2009; Wulff et al., 2009; Canolty et al., 2010; Nacher et al., 2013). The cross-frequency coupling may serve as a mechanism for the regulation of communications between different spatiotemporal scales (Palva et al., 2005, 2010; Holz et al., 2010). The phase coupling between theta and gamma oscillations, namely, the modulation of the gamma amplitude by theta phase is the most studied phenomena of phase coherence (Buzsáki et al., 1983, 2003; Bragin et al., 1995; Mormann et al., 2005; Canolty et al., 2006; Sirota et al., 2008; Tort et al., 2008; Sauseng et al., 2009; Scheffer-Teixeira et al., 2012; Schomburg et al., 2014; Musaeus et al., 2020).

Cognitive processes encompass different types of memory, and episodic memory is one of them. Episodic memory is defined as personal experienced events in the context of space and time (‘what happened to me where and when?’) (Tulving, 1985). The ability to simultaneously integrate these aspects of a unique experience is considered to be a feature of this type of memory (Griffiths et al., 1999; Pause et al., 2010). Interactions between the hippocampus and neocortical structures, such as the prefrontal and entorhinal areas, are believed to play a key role in this type of declarative memory (Chao et al., 2020). Besides, it has been shown that the hippocampus, a structure required for many types of memory, is anatomically and functionally connected with both the medial entorhinal cortex (mEC, where current sensory information is processed and the medial prefrontal cortex (mPFC), which helps direct neuronal information streams during intentional behavior (for review, see Colgin and Moser, 2010; Colgin, 2009, 2011; Mably and Colgin, 2018).

It was shown in experiments with rodents that these animals exhibited an episodic-like memory that resembles the episodic memory in humans, in which specific memory items are placed within a temporal and spatial context during encoding and retrieval (Tulving, 1983; Ennaceur and Delacour, 1988; Dupont et al., 2000). Based on the spontaneous preference for novelty by rodents, individual test object recognition tasks have been used to study episodic-like memory processes (Ennaceur and Delacour, 1988; Aggleton and Pearce, 2001; Fortin et al., 2002; Eacott and Norman, 2004; Langston and Wood, 2010), including the *what*, *where* and *when* components of personal experiences (Dere et al., 2005; Kart-Teke et al., 2006; DeVito and Eichenbaum, 2010).

There is evidence that different neural inputs carrying information from the overlying cortical structures converge to the hippocampus and create an image of the surrounding reality (Morris, 2001; Davachi, 2006; Witter et al., 2000; Eichenbaum et al., 2007). Thus, the significance of the hippocampus in episodic memory should depend on its ability to link these attributes of the unique experience into a spatiotemporal representation (MacDonald et al., 2011; Naya and Suzuki, 2011).

Episodic memory usually is violated during temporal lobe disturbances, especially temporal lobe epilepsy (Jokeit, Ebner, 1999; Helmstaedter, 2002; Hermann et al., 2006; Addis et al., 2007). Multiple investigations have shown that cell death in the hippocampus and related structures leads to a reorganization of circuits essential for encoding and retrieval, which may be the main cause of cognitive dysfunction in temporal lobe pathology (Glowinski, 1973; Jokeit and Ebner, 1999; Helmstaedter, 2002).

Recently, it has been suggested that theta/gamma coupling plays a critical role in episodic memory formation in the healthy brain and the lack of this coordination in the epileptogenic hippocampus (Shirvalkar et al. 2010; Lega et al. 2015, 2016). However, there is much to be learned and discussed: the oscillatory processes underlying the episodic memory and their disturbances in seizure pathology remain poorly understood. Whether prefrontal-hippocampal-entorhinal circuit functioning plays a specific role in this type of memory, and how this circuit alters during epileptogenesis are unknown yet. Thus, additional investigations are necessary to clarify this matter.

The main purpose of the present work is to investigate the characteristics of theta and gamma oscillations and theta/gamma coupling in the hippocampus and neocortical areas (mPFC and mEC) during execution of the episodic-like memory task by rats. We also tried to study changes in the rhythms and rhythm coordination after kainate (KA)-induced SE in periods free of interictal activity. More precisely, the theta power and the theta coherence between different structures and theta/gamma phase–amplitude coupling in all mentioned areas in healthy and epileptogenic rats were investigated. Using a kainate-induced SE model of epileptogenesis and recording local field potentials (LFPs), we demonstrate that the disruption of theta coherence and theta/gamma coupling in the prefrontal-hippocampal-entorhinal circuit in the brain of rats developing epilepsy (below referred to as epileptogenic rats) is linked to specific aspects of episodic-like memory deficits in these animals.

## Materials and Methods

The study was conducted in accordance with the ethical principles formulated in the Helsinki Declaration on the care and use of laboratory animals and the Regulations of the European Parliament (86/609/EC).

### Animals and surgery

Experiments were performed on young adult Wistar rats (males, 150-250 g, *n*=14) obtained from the Experimental Animal Center at the Institute of Theoretical and Experimental Biophysics (Puschino, Russia). Animals were housed in pairs under controlled conditions (22–24°C, 12 h light/dark cycle) with food and water ad libitum. They were assigned randomly to control or kainic acid (KA)-treated groups. The animals of the control group (n=8) were intraperitoneally injected with the saline, and the rats in the KA-treated group (n=6) were infused with kainate (i.p., usually they were injected once with 5 mg/kg of the drug. To identify the status epilepticus (SE) evoked by KA administration, the Racine Scale (Racine, 1972) was applied; stages 4-5 (tonic-clonic seizures, anticlockwise exploration with postural loss and falling) indicated the development of the SE. Animals that did not exhibit the SE were not included in the study. Control rats (the same weight and age, n=8) were injected with saline in an identical fashion and were treated similarly to animals in the KA group.

Prior to the beginning of experiments, animals were exposed to a surgery under general anesthesia (18 mg/kg Zoletil plus 12mg/kg xylazine, i.m.) in a stereotaxic apparatus (Kopf Instruments). The body temperature was maintained with a heating pad, and the cardio-pulmonary state during the surgery was monitored with a pulse oximeter (Oxy9Vet Plus, Bionet, South Korea). Depth recording electrodes (insulated nichrome, 0.05 mm diameter) were implanted using the brain atlas (Paxinos & Watson, 1998) into the hippocampus (field CA1: AP=-3.8, ML=2, DV=3), the medial prefrontal cortex (mPFC: AP=3, ML=0.2, DV=3), and the medial entorhinal cortex (mEC: AP=-8.5, ML=4.5, DV=6). The reference electrode was implanted into the occipital bone over the cerebellum. The whole complex was fixed on the head with dental acrylic resin. Animals were allowed to recover for one week, after which they were handled individually for 2 consecutive days for 40 min several times a day until the experiment was started.

### Electrophysiological records and analysis

The apparatus for testing the animals consisted of a square open field (80 x 80 x 50 cm) that was situated in an evenly illuminated room (15 lux) with ambient noise masked by a white-noise generator and several spatial cues. Visual cues were visible on the surrounding walls. Rat behavior was monitored with a video camera, and exploration of rats was analyzed off-line with a computer tracking system (EthoVision XT10). Local field potentials (LFPs) were recorded in behaving rats at the day time between 10 a.m. and 3 p.m. Recording was performed using a multichannel system (Model W8 Multi Channel Systems, Germany). Tree channels were recorded simultaneously, the sampling rate was 5 kHz. Paroxysmal activity (high-amplitude events or interictal spikes whose amplitude was three times higher than the standard deviation of the base signal) was eliminated from the general recording using a digital filter; then, the signals were summed, and the ratio of the obtained signal to the averaged values in the control was calculated. The analysis of phase relationships was carried out using the wavelet transform.

In the rhythmic activity, the ranges of theta (4-10 Hz), slow gamma (25-50 Hz), and fast gamma oscillations (55-100 Hz) were identified. The spectral power of these rhythms was measured in the animals in each exploration episode. Using the wavelet transform, the instantaneous phase and the amplitude of rhythmic activities were calculated for each signal with a step of 1 Hz in the range from 1 to 150 Hz. Coherence between theta oscillations in the hippocampus with theta rhythm in the mPFC and with those in the entorhinal cortex was calculated from the probabilistic distribution of the phase difference (with absolute coherence of theta rhythm in the two structures; the maximum of the probabilistic distribution of the phase difference is 1). In all structures, we also analyzed the coupling of gamma oscillations (slow and fast) with the phase of the hippocampal theta rhythm. Coupling was assessed as a measure of the heterogeneity of the distribution (Kullback–Leibler distance) of high-amplitude gamma events in the phase of theta cycle (strong coupling of rhythms was reflected in a high heterogeneity of distribution and in a high value of the modulation index).

### Object recognition task

To investigate the influence of KA-induced SE on cognitive brain functioning, the episodic-like “what-where-when” memory task was applied (Kart-Teke et al., 2006). In this memory tasks, we used the modified experimental paradigm described by de la Prida and colleagues (Inostroza et al., 2013b). Two sets of objects (4 objects each) differing in terms of height (10–15 cm), base diameter (8–10 cm), color, shape, and surface texture were used. At first, habituation sessions were performed (twice a day), which consisted of 10 min free open field exploration. After habituation, rats were tested for object–place recognition memory. Each trial consisted of two sample phases (3 min), followed by a test phase (3 min) after an intertrial period of 50 min. Episodic memory was assessed by comparison of the exploration time of various objects. Spatial coordinates were tracked simultaneously with the recording of LPPs, and the electrical activity of the hippocampus and cortical regions was compared with the animal’s exploratory behavior. In the episodic-like what-where-when memory task, the animals had to discriminate a novel object from familiar objects they saw in samples 1 (100 min ago, test 1) and 2 (50 min ago, test 2). A schematic illustration of the task is represented in Fig. 1. The objects were: A1 – the old stationary object, A2 – the old displaced object, B1 – the recent stationary, B2 – the recent displaced. Odor cues were removed after each trial by cleaning objects and the open field with alcohol 10%.

**Fig. 1.**
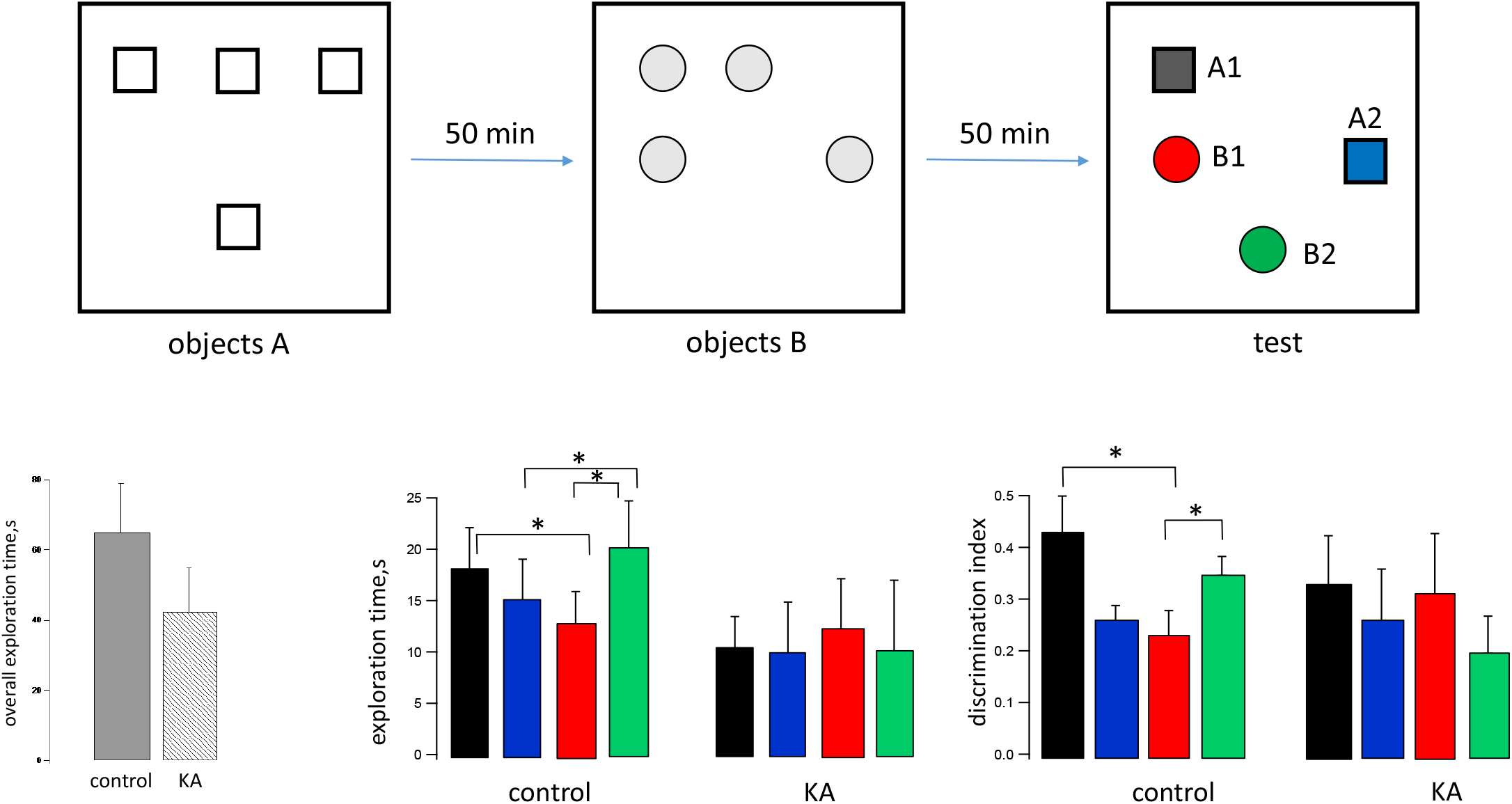
Episodic-like memory task. **A.** The episodic-like memory task consists of two sample phases and a test phase. In each sample phase, rats encountered two sets of four identical novel objects (old objects, sample 1; recent objects, sample 2). In the test phase, the objects were mixed together. Two of them were placed in the same location as in sample phases (A1, old stationary; B1, recent stationary). The other two objects were placed in new locations (A2, old displaced; B2, recent displaced). **B.** Total exploratory time for each animal group in the test phase. **C.** Distribution of exploration time per object in the test phase for the control and epileptic groups. A1, black, A2, blue, B1, red, B2, green. **D.** The mean discrimination index for objects. Data are represented as the mean ± SEM (n = 8 control, n = 6 KA-treated). \**p* ≤ 0.05.

### Behavioral data analysis

During the task, animals encountered two groups of objects in different locations and at different times. This task has been previously shown to test for nonverbal behavioral correlates of episodic memory (Pause et al., 2010). The rat was considered to explore an object, when it exhibited exploratory activity such as sniffing at least for 1 s, with its nose being directed at the object, and the central point of the body being within 1.5 cm from the object. The differences between the times spent for the exploration of different objects were examined to quantitatively evaluate the task performance. For each rat, the time spent for exploring the test object was converted to a discrimination ratio with the following formula: (Object_test_ – Object_control_) / (Object_test_ + Object_control_). A discrimination value of zero indicates no preference (chance level); a positive value indicates a preferential exploration of the test object, and a negative value indicates preferential exploration of the control object (Ennaceur and Delacour, 1988). For the episodic-like memory task, we defined the where memory index to quantify the time of the exploration of the object that had recently changed location as the proportion of the time spent for exploring a recent displaced object (B2) against a recent stationary one (B1) in a manner similar to that applied by DeVito and Eichenbaum (2010): Where = (B2 – B1) / (B2 +B1). The when memory index was defined as the proportion of time spent for exploring an old stationary object (A1) versus the time spent for the recent stationary object (B1) according to the following formula: When = (A1 – B1) / (A1 + B1). For these ratios, a value of zero indicates no preference (chance level), whereas a positive value indicates a preferential exploration of the recent displaced object for where and of the old stationary object for when. To evaluate against chance exploration of objects, we also defined discrimination indices (DIs) in the test phase of the episodic-like what-where-when task by dividing the time spent for exploring each of the objects by the total exploratory time on all four objects (chance level at 0.25) (see Fig. 1).

### Histology

One month after the kainate administration, morphological changes in the dorsal hippocampus were analyzed. After fixation in 4% paraformaldehyde (48h at 4°C) and cryoprotection in a gradient of sucrose (10% and 20% sucrose at 4°C for 24h each), brains were rapidly frozen in the vapor phase of liquid nitrogen and stored at −80°C. Coronal sections (15 μm) were cut with a cryostat at −19°C (Thermo Shandon Cryotome E, Thermo Scientific, USA) and collected on gelatin-coated slides for subsequent histochemical staining. Neurodegeneration was revealed by Nissl staining using a standard protocol. Slide-mounted sections were submerged in bidistilled water with acetate buffer for 5 min and stained in fresh 0.1% cresyl violet for 5–8 min until the desired depth of staining was achieved. The Nissl stained slides were then dehydrated through graded ethanols, cleared in xylene, and coverslipped with Eukitt (Fluka, Germany) mounting medium. Bright field images were acquired on a Nikon E200 microscope (Nikon, Japan) with a Sony alpha5000 (Japan) camera. All tissue sections were photographed under identical conditions. In the Nissl-stained sections of the right hippocampus, neuronal quantification was carried out in the dentate hilus, CA3 (counting frames 300×300 μm), and CA1 (counting frames 400×300 μm) fields. At least four different sections were evaluated from each animal. All analyses were carried out using the ImageJ software (1.43u, USA).

### 2.5. Statistical analysis

The results are presented as mean ± standard deviation. All statistical tests were performed using the IgorPro software (version 6.35, WaveMetrics, USA) and a library of online statistical calculators https://www.socscistatistics.com/. Normality was confirmed with the Kolmogorov–Smirnov test. In these cases, the data were analyzed with multivariate ANOVA with tasks and objects (discrimination ratios or exploration times) as the within-subject factors, and groups (control, epileptic) as the between-subject factor. Within-group comparisons were performed using the paired Student’s *t*-test. Statistical comparisons between the two groups were made using the unpaired *t*-test or the Mann–Whitney U test. Nonparametric statistics was used to avoid assumptions about (the) uniformity of variances or normal distributions; *p* < 0.05 was considered statistically significant.

## Results

Earlier, electrophysiological and histological experiments with healthy and kainate-treated rats during their behavior in the open field have been performed, and the results obtained in these experiments have been published elsewhere (Malkov et al. “Rhythmic Activity in the Hippocampus and Entorhinal Cortex is Impaired in a Model of Kainate Neurotoxicity in Rats in Free Behavior”. Neurosci. Behav. Physiol. 2021, 51, 73–84 (https://doi.org/10.1007/s11055-020-01041-7). The analysis of hippocampal LFPs recorded in the KA-treated animals revealed a paroxysmal activity (high-amplitude events or interictal spikes) and alterations in the theta and gamma oscillations in the hippocampus and the mEC. Post mortem histological analysis in animals of this group identified significant damage to the hippocampus and dentate gyrus. However, no significant differences in exploratory behavior in the open field were found in the rats of these two groups.

LFP records showed that the theta rhythm was dominant in the hippocampus, mEC and mPFC during the tests. Analysis of oscillation parameters in the hippocampus showed that movement of the rats from the periphery to the center of the field was accompanied by a significant increase in the frequency of the theta rhythm (7.46 ± 0.4 Hz for exploration vs. 6.81 ± 0.14 Hz for background, F(1,10) = 6.913, *p* = 0.0465; see results in Malkov et al., 2021). The slow gamma rhythm showed no significant changes as the rats started exploration. However, the fast gamma rhythm underwent significant changes: its frequency decreased significantly when the rats moved from the peripheral to central zones (from 64.0 ± 1.2 to 60.5 ± 1.85 Hz, p < 0.05, F =10.924, *p* = 0.02); its power level did not change.

When the KA-treated animals moved from the periphery to the center of the field for exploration, the theta rhythm power and frequency did not change in the hippocampus; there was also no significant change in slow and fast gamma oscillations (the frequency of fast gamma oscillations decreased in controls). However, it should be noted that in these rats fast gamma oscillations initially had significantly higher frequency than those in healthy rats (66.0 ± 2.0 Hz vs. 60.5 ± 1.8 Hz, F = 7.23, p = 0.025). The frequency of the slow gamma rhythm did not change when animals moved from the periphery to the center of the field, though control rats showed a minor increase in frequency.

Afterwards the rats of these two groups were tested in the paradigm of episodic memory with simultaneous recording of LFPs. Here we describe the results obtained in these experiments.

### Characteristics of what, where, when memories in control and epileptogenic animals

#### Control animals

At first, rats were examined using the single-trial object recognition tasks at 50 min intervals between the sample 1 and sample 2 episodes. Then, after 50 min, the rats were subjected to a test phase, in which the objects were mixed and rats were tested for binding of the *what-where-when* memory triad (Fig. 1). In the test phase, healthy animals exhibited differential exploration, spending more time searching for an old familiar object A1 than a recent familiar object B1 that remained stationary (Fig. 1C, left, A1 black 23.65 ± 4.1 s vs B1 red 12.6 ± 3.2 s; F(1.10) = 5.61, *p* = 0.039). They also showed biased exploration, spending more time searching for a recent displaced object B2 than the stationary object B1 (Fig. 1C, left, B2 green 26.3± 4.7 s vs B1 red 12.6 ± 3.2 s; F(1.12) = 5.22, *p* = 0.041). Thus, the control animals preferred to explore the old stationary object (A1) and the recent displaced object (B2); the discrimination indices for these objects were 43 ± 7% and 35 ± 4%, respectively (Fig. 1D, left). The normal animals also distinguished with a high probability the object displacement (indicator *where* was 0.33 ± 0.12) and their temporal order (indicator *when* was 0.35 ± 0.16) (accidental exploration of the presented objects, without the involvement of episodic memory, would result in *when-where* parameters of around zero).

#### KA-treated animals

A month after the KA injection, rats spent a shorter time for exploring the presented objects compared to the control animals (Fig. 1B). Objects A1 and B2, which were the most interesting for healthy animals, were studied with less attention and, obviously, were not remembered by KA-treated rats, since they spent almost the same time studying them and other objects. The discrimination indices for A1 and B2 were 33 ± 10% and 20 ± 7%, respectively (Fig. 1D, right). The *where* memory index (discrimination ratio) was 0.00 ± 0.17, and the *when* memory index (discrimination ratio) was 0.05±0.18. In general, the epileptogenic animals did not have preferred objects for research.

### Changes in the oscillatory activity during the episodic-like memory test

#### Control animals

In the hippocampus, the power of theta and slow gamma rhythms did not change when rats explored various objects (Figs. 2, 3A, B) Instead, the power of fast gamma oscillations was significantly lower when animals studied displaced objects (A2 0.023 ± 0.004 a.u., and B2 0.024 ± 0.005 a.u.) compared to that when they investigated stationary ones (A1 0.046 ± 0.013 a.u., and B1 0.026 ± 0.004 a.u.) F(1.24) = 5.89, *p* = 0.023), *p* = 0.023) (Fig. 3C). The difference in the power of fast gamma oscillations was especially pronounced when rats explored A1 and A2 (old stationary and old displaced objects) (F(1.10) = 6.12, *p* = 0.033) (Fig. 2). Thus, the main factor that induced changes in the fast gamma rhythm was the displacement of the objects.

**Fig. 2.**
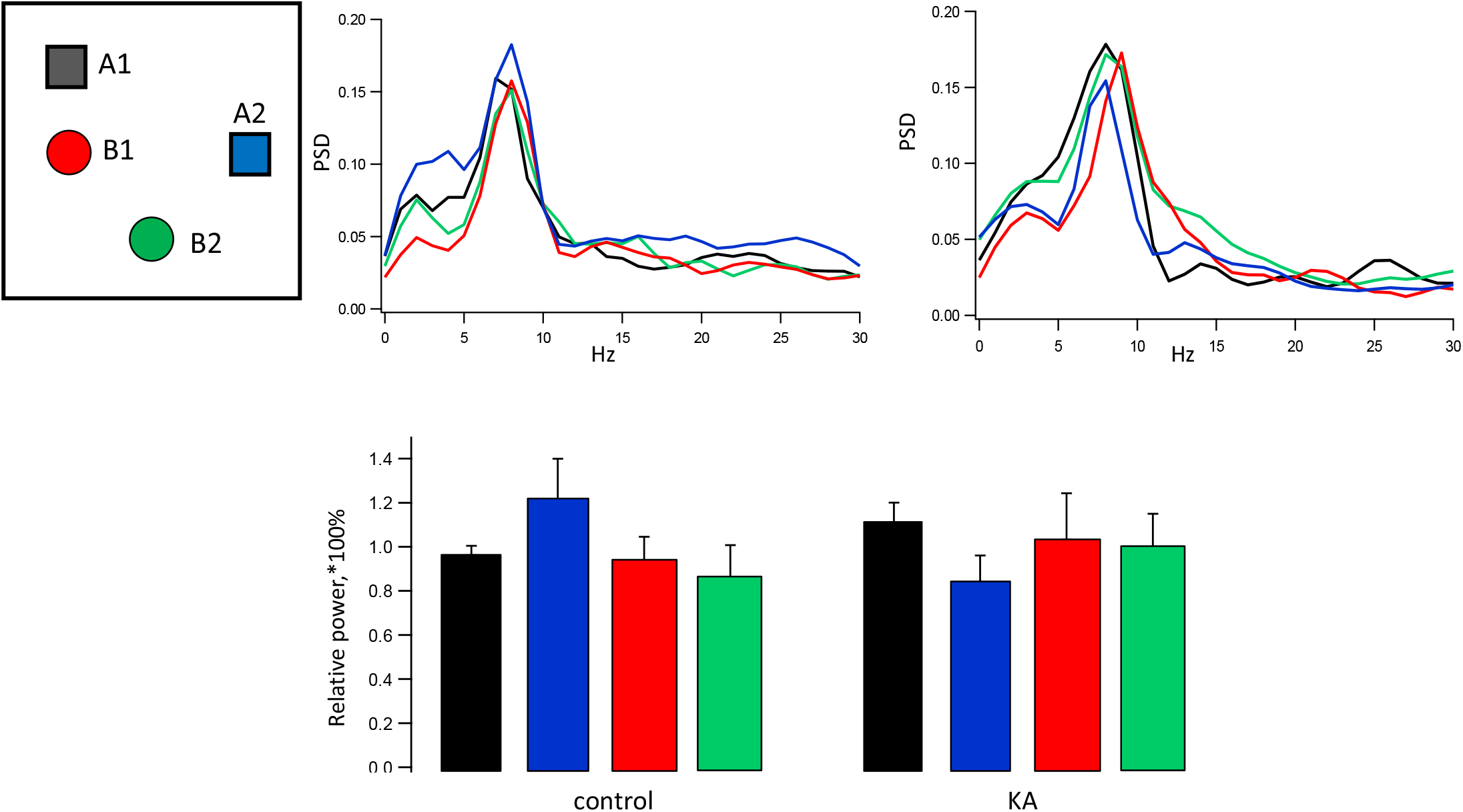
Hippocampal theta activity in the episodic memory test. **A.** Averaged power spectral density diagrams of the theta-band LFP during exploration of various objects. A1 black, A2 blue, B1 red, B2 green. **B.** Mean values of the theta band power per object normalized to the exploratory behavior level. Data are represented as the mean ± SEM (n = 8 control, n = 6 epileptic). \**p* ≤ 0.05.

**Fig. 3.**
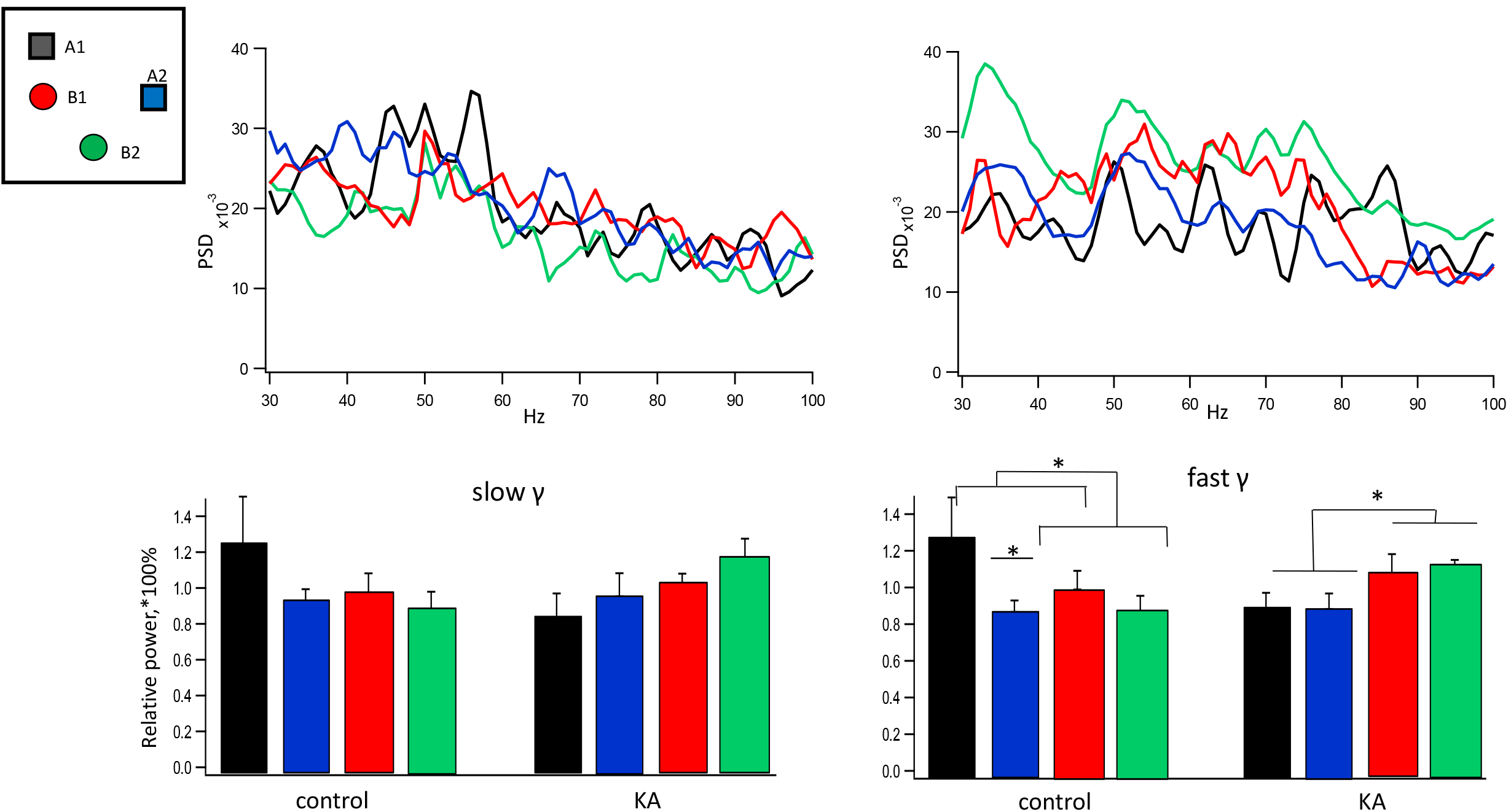
Hippocampal gamma activity in the episodic memory test. **A.** Averaged power spectral density diagrams of the gamma-band LFP during exploration of various objects. A1 black, A2 blue, B1 red, B2 green. **B.** Mean values of the slow (on the left) and fast (on the right) gamma band power per object normalized to the exploratory behavior level. Data are represented as the mean ± SEM (n = 8 control, n = 6 KA-treated). \**p* ≤ 0.05.

In the mPFC and mEC, as in the hippocampus, only the fast gamma rhythm changed when rats explored various objects. These changes were similar to the hippocampal ones: fast gamma oscillations were significantly lower when animals explored displaced objects (A2 and B2) compared to the oscillations recorded when they explored stationary objects (A1 and B1) (mPFC F(1.24) = 4.47, *p* = 0.045; mEC F(1.24) = 4.38, *p* = 0.047).

#### KA-treated animals

In the hippocampus, the power of oscillations during the exploration of different objects changed in the fast gamma range only, as was the case in the control animals. But in this group, the main modifying factor was the object temporal order (old or recent item), but not its placement. Namely, during the exploration of the recent objects (B1 and B2) the fast gamma power was higher (B1 0.076 ± 0.027 a.u.; B2 0.084 ± 0.022 a.u.) than during examination of the old ones (A1 0.064±0.014 a.u.; A2 0.059± 0.009 a.u.), F(1.16) = 7.8, *p* = 0.013) (Figs. 2, 3).

In the mPFC and mEC, as in the hippocampus, only fast gamma rhythm changed when rats explored various objects. The changes were similar to the hippocampal ones: during the exploration of the recent objects (B1 and B2), the fast gamma power was higher when rats explored the old ones (A1 and A2) (F(1.16) = 7.82, *p* = 0.013.

### Changes in the interstructural theta coherence

Here, we investigated the coherence between the hippocampal theta rhythm and theta oscillations in the mPFC and those in the mEC.

#### In control animals

the theta coherence between the hippocampus and the mPFC did not change when rats explored various objects (Fig 4A, C). Instead, the coherence between the hippocampus and mEC was significantly higher when rats explored the recent objects (B1 0.43±0.04 a.u.; B2 0.36±0.034 a.u.) than during the exploration of the old ones (A1 0.31 ± 0.04 a.u.; A2 0.37 ± 0.05 a.u.), *p* < 0.05 (Fig 4 B, D).

**Fig. 4.**
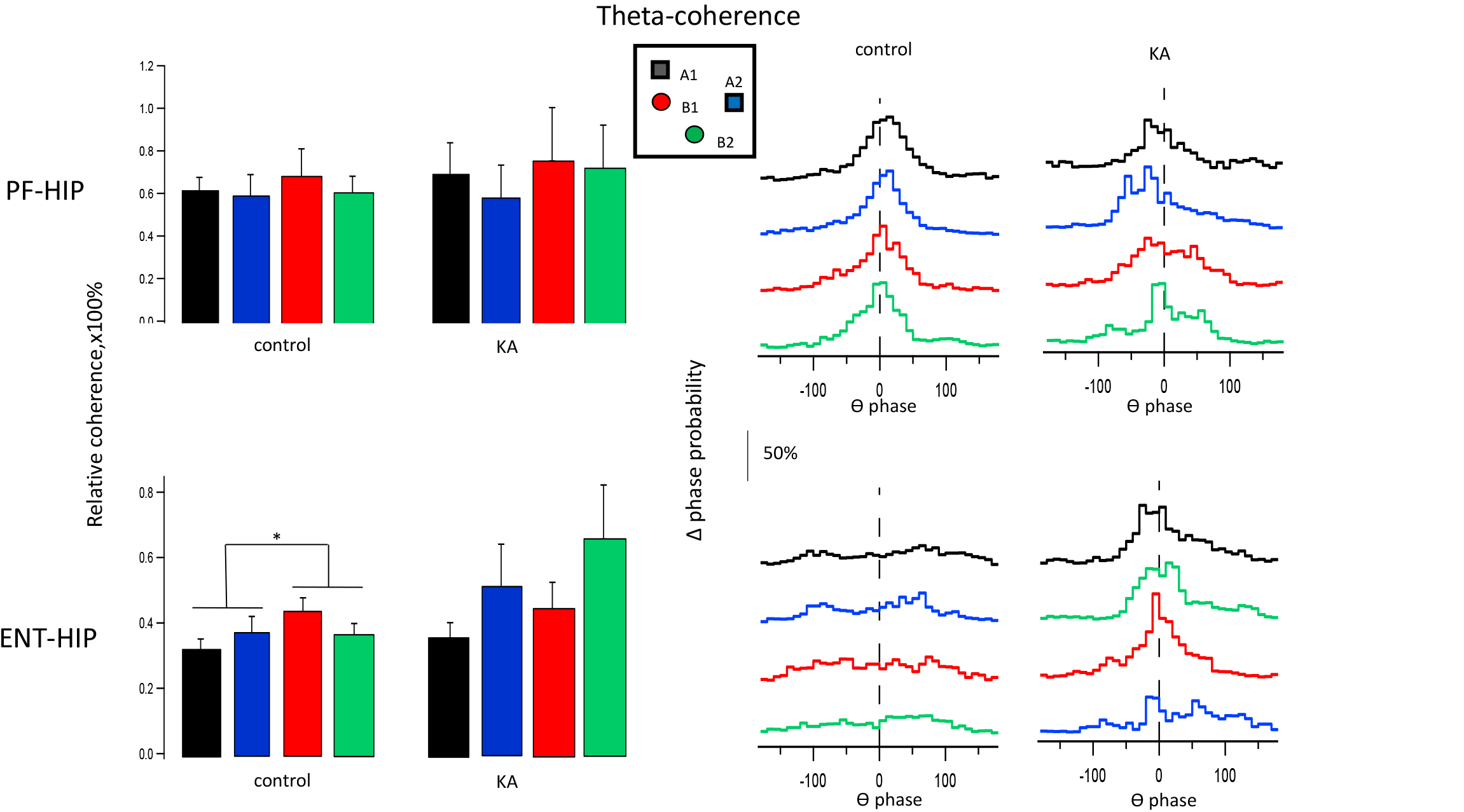
Interstructural theta coherence. **A, B**. The relative theta coherence per object between the hippocampus and mPFC **(A)**, and between the hippocampus and mEC **(B)**. Data are represented as the mean ± SEM, \**p* ≤ 0.05. **C, D.** Distribution of the probabilities of the phase difference of theta activity in the pair of hippocampus–mPFC (**C**) and hippocampus–mEC (**D**). A1 black, A2 blue, B1 red, B2 green.

#### In KA-treated epileptogenic animals

no differences in theta coherence between the hippocampus and cortical regions were found when they examined different objects (Fig 4). The mean level of coherence between mEC and hippocampus during object exploration was by 30% higher (0.48±0.08 a.u. vs 0.37±0.04 a.u., F(1.11) = 8.0, *p* = 0.016). No difference was observed in the mean level of coherence between mPFC and hippocampus during object exploration compared to the control.

### Modulation of local rhythms by the hippocampal theta phase (theta/gamma phase – amplitude coupling)

Here, we investigated how the phase of the hippocampal theta rhythm modulates the amplitude of the slow and fast gamma oscillations in all three structures, the hippocampus, the mPFC, and the mEC, in control and KA-treated rats. First, the theta/gamma modulation when rats performed the task was compared to its background level when animals moved in the same area of the open field but without objects (during habituation session). Then, relative changes in the theta/gamma modulation during exploration of different objects by rats were measured and expressed as a percentage of the mean exploration level.

#### Control animals

In our experiments, slow and fast gamma rhythms in the hippocampus of control rats were nested in the ascending and descending phases of the theta cycle, respectively, which agrees with the earlier data (Buzsáki et al., 1983; Bragin et al., 1995; Belluscio et al., 2012). The mPFC and mEC also demonstrated sufficient coupling of gamma activity with local theta oscillations. While in the mPFC the coupling pattern was similar to the hippocampal one, the gamma oscillations in the mEC were coupled to the lower part of the descending theta wave. In all structures examined, the exploration of objects was accompanied by an essential increase in the theta/gamma modulation, both in slow and fast gamma bands (Fig.5).

In the hippocampus, the theta/gamma modulation index increased during exploration from 6.56 ± 0.44 to 17.79 ± 2.8 (F(1.14) = 13.8, *p* = 0.0075) for slow gamma, and from 6.1 ± 0.7 to 18.3 ± 2.6 (F(1.14) = 27.5, *p* = 0.0012) for fast gamma band (Fig. 5). The theta/slow gamma coupling did not differ when rats explored various objects. On the other hand, the modulation index for theta/fast gamma coupling was higher near the stationary objects (A1: 112±19% and B1: 112±14%) compared to displaced ones (A2: 72 ± 8% and B2: 99 ± 10%; F(1.24) = 5.28, *p* = 0.013; Fig. 6).

**Fig. 5.**
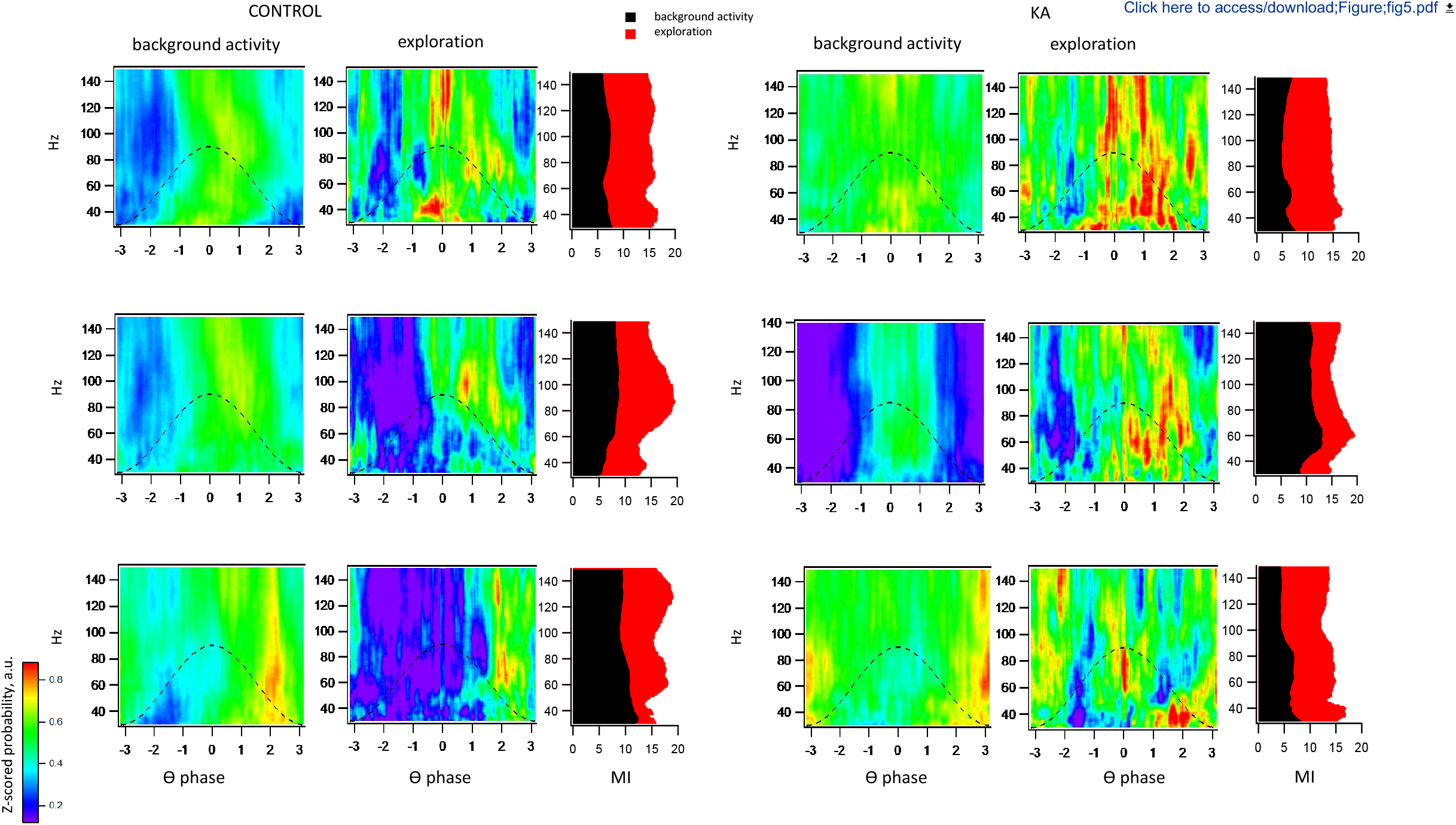
Changes in theta/gamma coupling during the transition of rats to exploratory behavior. **A, C.** Color-coded gamma power as a function of theta phase in the hippocampus (top), mPFC (middle), and mEC (bottom) in control (**A**) and KA-treated rats (**C**). **B, D**. Strength of theta modulation of gamma activity (bottom axis – modulation index) as a function of power frequency (left axis) in the mPFC, the hippocampus and MEC, respectively. Averaged data are represented.

**Fig. 6.**
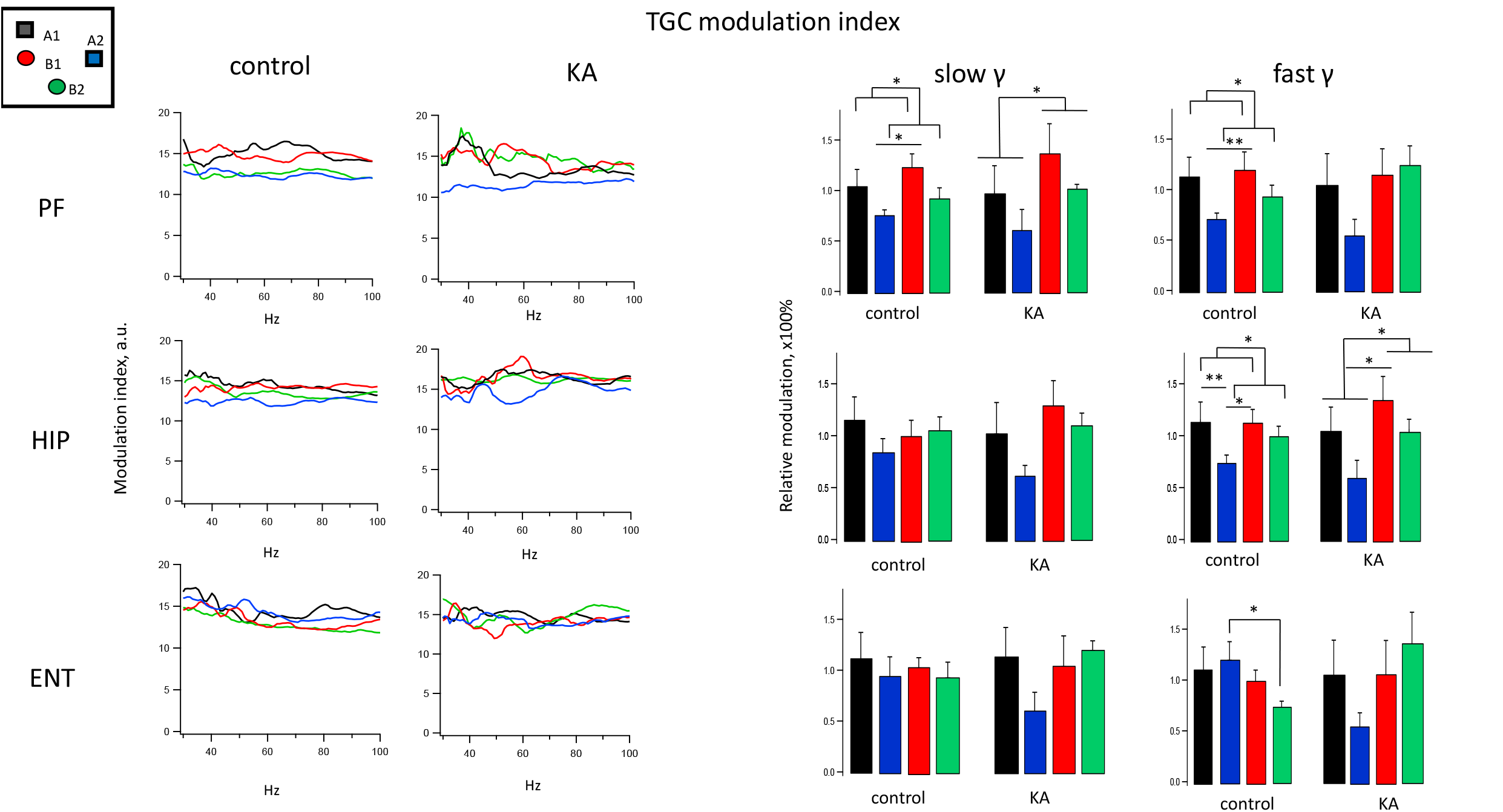
Theta-gamma coupling during the episodic memory test. **A.** Strength of theta modulation of gamma activity (left axis – modulation index) as a function of power frequency (bottom axis) in the hippocampus (top), mPFC (middle) and MEC (bottom) during the exploration of objects by rats. **B.** The normalized modulation index for slow and fast gamma rhythms during the episodic memory test when rats explored objects (A1 black, A2 blue, B1 red, B2 green). Data are represented as the mean ± SEM (n = 8 control, n = 6 KA-treated). \**p* ≤ 0.05.

In the mEC, when animals explored the objects, the theta/slow gamma modulation significantly increased, from 11.5 ± 3.13 to 18.6 ± 2.3 (F(1.14) = 13.8, *p* = 0.0075); vice versa, the theta/fast gamma modulation only slightly increased, from 12.8 ± 3.4 to 19.0 ± 1.8 (F(1.14) = 3.35, *p* = 0.1) (Fig. 5). Theta/slow gamma coupling did not differ when rats explored various objects (just like in the hippocampus). Conversely, theta/fast gamma coupling was higher, when animals explored old displaced object (A2: 120±18%) compared to recent displaced one (B2: 73±6%; F(1.10) = 5.55, *p* = 0.04; Fig. 6).

In the mPFC, as well as in hippocampus and mEC, theta/gamma modulation was higher when animals explored objects (for slow gamma 17±2.36, F(1.14) = 13.1, *p* = 0.0084; for fast gamma 15.3 ±2.2, F(1.14) = 16.3, *p* = 0.005) compared to the background activity (7.9±0.58 and 7.5 ±0.85, respectively). Theta/slow gamma coupling was significantly higher when rats explored stationary objects (A1 104±17% and B1 122±14%) compared with those when they studied displaced ones (A2: 74±6% and B2: 94±3%; (F(1.24) = 5.89, *p* = 0.03; Fig. 6). The same was observed in relation to theta/fast gamma coupling (A1 112±20%, B1 119±18%, A2 69±7% and B2 92±12%; F(1.24) = 5.32, *p* = 0.023; Fig. 6).

#### KA-treated rats

In KA-treated SE animals, phase preference of the hippocampal and cortical oscillations to the phase of the hippocampal theta cycle was the same as in the control group. However, it should be noted that, during background activity, the coupling of fast gamma waves to the phase of the theta cycle was significantly higher in the KA group compared to the control one (F(1.12) =17.1, p = 0.0016).

In the hippocampus, the exploratory behavior caused a notable but statistically insignificant increase in the theta/gamma coupling from 10.1±2.36 to 16.92±3.1 for slow gamma, and from 15.6±2.7 to 19.1±4.1 for the fast gamma band (Fig. 5). Theta/slow gamma coupling did not differ when rats explored various objects (as was the case in control rats). (A1 102±30%, B1 124±28%, A2 60±12%, and B2 101±5%, Fig 6). However, theta/fast gamma coupling, in contrast to the control animals, was significantly higher when the rats explored the recently presented objects (B1-134±30%, and B2-103±13%) compared to the old ones (A1 96 ±30%, and A2 59 ± 13%; F(1.16) = 6.53, *p* = 0.021; Fig. 6).

In the mPFC, an insignificant increase in the modulation index during the transition from background to exploratory activity was also observed, from 7.0 ±1.1 to 17.2 ±4.4 for slow gamma, and from 6±0.83 to 13.5 ±3.4 for the fast gamma band. Theta/slow gamma coupling was higher when animals explored the recently presented objects (B1 135±31% and B2 101±5%), compared to the old ones (A1 97 ±28% and A2 59 ± 22%; F(1.16) = 5.99, *p* = 0.026). Theta/fast gamma coupling did not differ when rats explored various objects (Fig. 6).

In the mEC, the theta/gamma modulation increased when animals explored the objects from 7.8±1.8 to 17.6 ±3.1 for slow gamma and from 6.7±1.1 to 16.5 ±3.3 for the fast gamma band, but the changes were statistically insignificant. Theta/slow and theta/fast gamma coupling did not change when rats explored various objects (Fig. 6), but minor alterations in the modulation pattern were observed compared to the control.

Thus, in KA-treated rats, the modulation of local rhythms by the hippocampal theta phase was disturbed compared to that in control animals.

### Negative correlation of the theta/gamma coupling with the success of execution of the behavioral episodic-like memory test

In the control group, the degree of discrimination of objects negatively correlated with theta/gamma modulation in all structures; i.e., the better the animal explored the object, the lower was the theta/gamma coupling. Besides, successful discrimination of objects displacement negatively correlated with theta/fast gamma coupling in the hippocampus: i.e., the better the animal executed this task, the lower was the hippocampal theta/fast gamma coupling.

On the whole, most of the parameters measured during the exploration of the objects with novel properties (earlier presentation or displacement) negatively correlated with the success of their discrimination. It was assumed that the successful recognition of the temporal order of the objects and their placement depended not on the strength of theta/gamma coupling *per se*, but on the difference in this strength during the exploration of the objects with diverse properties. We analyzed the correlation between the success in performing the *when/where* tasks and the difference in the modulation index of slow and fast gamma oscillations (ΔMI (A1-B1) and ΔMI (B1-B2), respectively. As a result, we were able to show a pronounced correlation between the theta/slow gamma and theta/fast gamma modulation in the hippocampus and the success in the execution of the *when* task (Pearson’s coefficient V_Pr = 0.865 and V_Pr = 0.83, respectively, *p*<0.05) (Fig. 7). We also showed a correlation between the success in performing this task and theta/slow gamma as well as theta/fast gamma modulation in the mPFC (Pearson’s coefficient V_Pr = 0.8 and V_Pr = 0.77 respectively, *p*<0.05) (Fig. 7). In the mEC, the performance of the *when* task correlated with theta/slow gamma (V_Pr = 0.78, *p*<0.05) and theta/fast gamma coupling (V_Pr = 0.76, *p*<0.05).

**Fig. 7.**
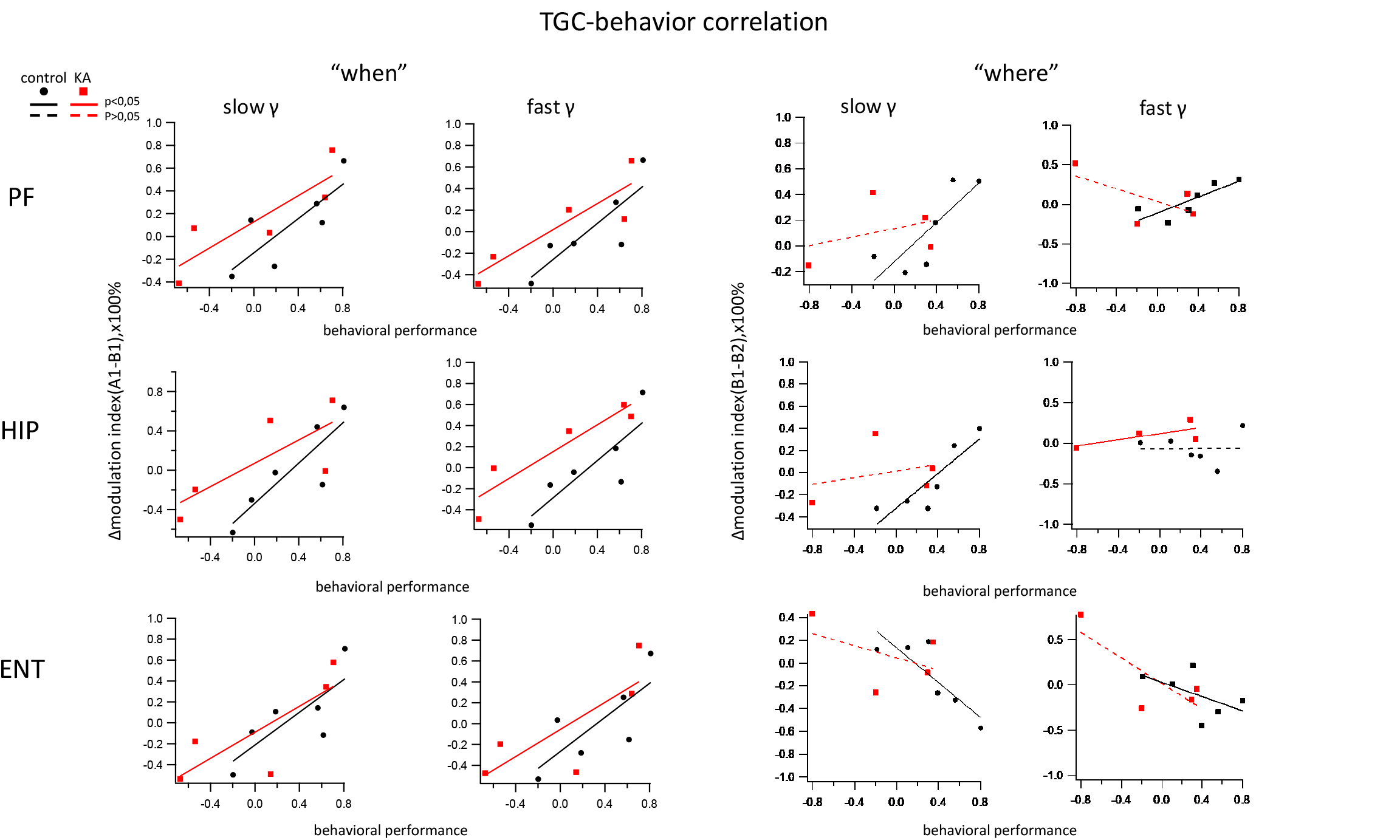
Correlation of electrophysiological parameters and the behavioral performance in control (black) and KA (red) groups. **A**. Correlation between a shift in theta/gamma modulation around objects A1 and B1, and the discrimination index *when*. **B**. Correlation between a shift in theta/gamma modulation around objects B1 and B2, and the discrimination index *where*. Solid lines denote the statistical significance of Pearson correlation (*p* < 0.05), dotted lines are for insignificant (low) correlation.

In the case of the *where* task, theta/gamma coupling also correlated with ΔMI in the mPFC (V_Pr = 0.82 for slow and V_Pr = 0.81 for fast gamma, *p*<0.05). In the hippocampus, this dependence was significant for slow gamma band only (V_Pr = 0.87, *p* <0.05). In the mEC, the performance of the *where* task negatively correlated with the chosen parameter ΔMI (B1-B2) (V_Pr = −0.85 for the slow gamma and V_Pr = −0.79 for the fast gamma, *p*<0.05), indicating that only in the mEC the theta/gamma coupling near the displaced object was higher than near the stationary object.

In the epileptogenic animals, although the performance score in the *when* task correlated with ΔMI (A1-B1) to the same degree as it was in control animals, drastic violations were observed during the performance of the *where* task. Only theta/fast gamma coupling substantially correlated with the *where* score (V_Pr = 0.68, *p*<0.05). For hippocampal fast gamma band, as well as slow and fast gamma bands in the mPFC and mEC, the correlation was insignificant.

## Discussion

Overall, our findings provide new insights into the neural mechanisms of the episodic-like memory and their impairment in kainate-treated SE rats.

### SE rats were epileptogenic in our experiments

Previously it was shown that the long-lasting SE can lead to the development of chronic epilepsy with recurrent spontaneous seizures (Lothman et al., 1990; Hellier et al. 1998; Bragin et al. 1999). Although we were unable to record spontaneous seizures in SE rats, however, high-amplitude interictal spikes were revealed in the hippocampal LFPs of these animals; these were never observed in the control rats. Earlier it was also shown that the absence of spontaneous seizures in SE animals is observed only in behavior, while paroxysmal activity is clearly recorded in the EEG (Mazarati et al., 2002). Authors found a statistically significant correlation between the frequency of acute events (spikes) during SE and those during the interictal period. Besides, interictal spikes may correlate with the degree of hippocampal injury in epilepsy patients (Spencer et al., 1992). Moreover, a transition from SE to interictal spiking in a rodent model of mesial temporal epilepsy (Wang et al., 2019) and a transition of interictal discharges to the onset of ictal activity, a phenomenon termed the interictal-to-ictal transition, have been shown (Gotman, Marciani, 1985; Lange et al., 1983; Gotman, 1991). In our work, we have not observed spontaneous seizures, probably because one month is the term not sufficiently long for their distinct manifestation: they usually occur on the average 2.5 months after kainate-induced SE (Hellier et al., 1998). On the other hand, the electrophysiological changes we observed might be the early biomarkers of degenerative processes developing during epileptogenesis (Betjemann, Lowenstein, 2015; Lin et al., 2019). Indeed, we revealed a significant decrease in the number of neurons in the hippocampus and dentate gyrus in SE rats. Thus, we believe that our experimental animals injected with kainate were at an early stage of the development of temporal lobe epilepsy.

### Control but not epileptogenic animals recognized objects and remembered their place and order of presentation during episodic-like memory task

We revealed that, during the *what-where-when* episodic-like memory test, healthy animals recognized objects explored during separate trials and remembered their place and the temporal order. Namely, they spent more time exploring an old familiar object than a recent familiar one that remained stationary; they also differently explored stationary and displaced objects, spending more time near a recent displaced object than near the stationary one. Thus, the animals preferred to explore the old stationary and the recent displaced objects. These data are consistent with the findings of previous experiments on normal rats (Kart-Teke et al., 2006; Inostroza et al., 2013b).

One month after SE, the rats exhibited an impairment of episodic memory for the *what-where-when* triad. Unlike healthy rats, epileptogenic animals did not recognize objects during the test. The lack of preference in the exploration of the objects that changed their location or the temporal order, which obviously was observed in the normal animals, indicates a significant impairment of episodic-like memory in epileptogenic rats as tested in the *what-where-when* coupled task. Earlier it has been demonstrated that this impairment in kainate-treated epileptic rats was specific for the integrated *what-where-when* memory only, whereas the rats did not exhibit any cognitive deficit if the individual components were tested in separate tasks not requiring any binding of spatiotemporal context with an event (Kart-Teke et al., 2006; Inostroza et al., 2013b). Previously, de la Prida with colleagues showed that epileptic rats were able to solve a complex version of an object recognition task with a 5 min retention interval (Suarez et al., 2012) and the allocentric version of the water maze (Inostroza et al., 2011); all these facts indicate that they were able to manage high cognitive loads. Therefore, failure in the *what-where-when* task points to specific dysfunction of epileptic rats to integrate spatial and temporal aspects of events when tested at once (Chao et al., 2020, for review).

### Specific changes in the fast gamma rhythm during object exploration. Alterations in hippocampal and neocortical oscillations in epileptogenic rats

In our experiments, we were able to compare the power of oscillations measured during the exploration of objects by rats in the episodic-like memory test with the background (or basic) rhythm parameters measured when rats explored the open field without objects.

Interestingly, in our work we revealed that only fast gamma rhythm changed significantly when rats explored different objects. Namely, the power of fast gamma oscillations was significantly higher when animals explored the stationary objects (old A1 and recent B1) compared with its mean basic value, and it was lower during exploration of displaced items (A2 and B2), i.e., objects with associative novelty, presented in updated spatial locations. We found that the difference in the power of fast gamma was especially pronounced when rats explored the old familiar stationary object (A1) compared with an old familiar displaced one (A2) (Fig. 3). Earlier, it was demonstrated that hippocampal fast gamma dominates during exploration of novel object–place pairings (Bieri et al., 2014; Zheng et al., 2016). In contrast, in our experiments, fast gamma activity significantly increased during the exploration of objects without obvious novelty properties. In the study by Zheng with co-authors, another behavioral testing paradigm was used, with shorter intertrial intervals (10 min), presumably sufficient for capturing the novelty encoding. In our case, with larger timescales (intertrial intervals of 50 min), the presentation of previously explored objects could lead to prevalence of reactivation memory processes rather than new encoding. Thus, one possible explanation of our results is that the increase in the fast gamma rhythm reflects the reactivation of memory about familiar context, as it was shown by Cammarota with colleagues (Radiske et al., 2017) for the rats in the fear-motivated avoidance paradigm. They revealed that the exploration of familiar objects can be accompanied by an increase in the power of hippocampal fast gamma oscillations; probably, this indicated the reconsolidation of memories of the object and its spatiotemporal context. Although in our present experiments, another experimental paradigm was used, we believe, that the same is true for estimation of the role of the fast gamma rhythm in the episodic-like memory task. Now then, it can be assumed that our results indicate the involvement of fast gamma oscillations in reactivation and updating (reconsolidation), at least, of the *where* component of episodic-like memory in rats.

We also demonstrated that, in all structures under investigation, the shift in fast gamma power was unidirectional during the exploration of objects by rats. Since the hippocampus, a structure required for many types of memory, has been shown to be connected anatomically and functionally to both mEC and mPFC (Colgin and Moser, 2010; Colgin, 2009, 2011; Mably and Colgin, 2018, for review), it is not surprising that the shift in the power of fast gamma rhythm in these structures when rats explored objects was unidirectional. Our findings also support the hypothesis that slow and fast gamma oscillations are functionally distinct in the hippocampal–mPFC–mEC network, as has been suggested previously for the entorhinal–hippocampal network (Colgin et al., 2009).

In SE animals, as in control rats, the power of oscillations during the exploration of different objects changed only in the fast gamma range; but in this group, the modifying factor was the order of the presentation of the object (*when* component), but not its placement, as it was in healthy rats (*where* component). Thus, our data indicate that, in epileptogenic animals, the contribution of the fast gamma rhythm to the recognition and encoding of the object placement in the episodic-like memory test. The impairment of the specific role of the fast gamma rhythm can disrupt the correct encoding of the place and sequence of events in personal memory.

As for the theta rhythm, our results are partially consistent with the previous data (Inostroza et al., 2013b). In this study, the power of theta recorded at CA1 stratum lacunosum-moleculare was disturbed when rats explored the spatial component (*where*) only in the epileptic, but not in control animals. According to the data of these authors, we did not observe significant changes in the power of theta oscillations recorded in the CA1 pyramidal layer in control rats during the exploration of objects, but we did not find any changes in the theta power in epileptogenic animals too.

### Theta coherence between the hippocampus and mEC increased in the control but not epileptogenic animals during the episodic-like memory task

Theta oscillations are known to be the dominant activity in the hippocampus, a structure that is critical for episodic memory (Aggleton et al., 1999; Steinvorth et al., 2005; Fortin et al., 2004; Ergorul and Eichenbaum, 2004; Day et al., 2003). Relative to the temporal organization of episodic memory, the hippocampus is essential to remembering unique sequences of events as well as the ability to disambiguate sequences (Fortin et al., 2002; Kesner et al., 2002; Agster et al., 2002; Kumaran and Maguire, 2006; Ross et al., 2009; Lehn et al., 2009; Brown et al., 2010; Tubridy and Davachi, 2011). The theta rhythm is thought to play a role in coordinating the activities during memory encoding (Winson et al., 1978; Pavlides et al., 1988). In this aspect, theta oscillations are involved in facilitating the information transfer from one brain region to another during sensory information processing. In agreement with this, we observed significant changes in the theta coherence between the hippocampus and mEC in control rats during the exploration of objects. Despite the lower level of coherence, the data dispersion was minimal, which allowed us to find significant coherence difference when animals explored the objects. The coherence was significantly higher during the exploration of recent objects by rats than when they explored the old ones, i.e., during activation of the *when* component of episodic-like memory. In contrast, in the epileptogenic animals along with an increase in the mean basic coherence level, there was no change in theta coherence between the hippocampus and mEC, when rats explored the objects.

Studies by other authors showed the coordinated theta oscillations across the hippocampal–entorhinal system (Buzsáki, 2002; Lubenov, and Siapas, 2009; Patel et al., 2012) and the coding of time and space by the theta rhythm, which can account for episodic memory retrieval (Hasselmo, 2009; 2012). These processes could underlie the increase in the theta coherence between the hippocampus and mEC during the exploration of objects with temporal order novelty that we observed in the rats in our experiments. Hyperactivation caused by kainate injection may lead to an increase in the level of hippocampal-entorhinal phasic interactions (Froriep et al. 2012, Kitchigina, 2018), followed by an increase in the background interstructural coherence (Malkov et al. 2021).

The impaired coordination of theta activity between the hippocampus and mEC during signal processing, which we found in epileptogenic animals, can disrupt signal coding and memory formation, in particular in terms of the temporal order of signals.

### Objects with signs of novelty caused different changes in theta/gamma coupling in healthy and epileptogenic animals

In experiments with the control animals, we showed that theta/slow gamma coupling in the mEC and theta/fast gamma coupling in the hippocampus and mPFC in rats exploring the stationary objects (A1 and B1) were stronger than in animals examining the replaced ones, i.e., objects with signs of spatial novelty (A2 and B2).

In the epileptogenic rats, in contrast to healthy animals, the theta/gamma coupling in the hippocampus and mPFC was stronger when rats explored the recently presented objects (B1 and B2), than during the exploration of old ones (A1 and A2). There were no changes in theta/gamma coupling in the mEC. Thus, it can be suggested that in epileptogenic rats, the objects with signs of temporal-order novelty (recency) caused an increase in theta/gamma coupling, an event different from that observed in the control.

Earlier, it has been proposed that, in the mEC, the coupling between theta phase and slow gamma oscillations facilitates the retrieval of previously learned tasks (Mormann et al., 2005; Tort et al., 2009; Shirvalkar et al., 2010; Bieri et al., 2014), although other works have suggested that the encoding rather than memory retrieval is associated with slow gamma modulation (Kemere et al., 2013; Trimper et al., 2014). At the same time, there is a hypothesis that theta-modulated fast gamma contributes to the encoding of current information (De Almeida et al., 2012; Newman et al., 2013; Cabral et al., 2014; Bieri et al., 2014). However, a more recent study has challenged the hypothesis that theta-modulated fast gamma is involved in the coding of current signals (Yamamoto et al., 2014).

The interaction of the theta rhythm and fast gamma oscillations in the hippocampus can play a key role in coordinating interactions between encoding and retrieval during exploration. Cammarota with colleagues (Rediske et al., 2017) showed that habituation to the environment before testing caused an increase in the theta/gamma coupling during further memory reactivation; this is obliquely consistent with our data. The authors declared that the strength of cross-frequency coupling in the hippocampus is an electrophysiological correlate of memory reconsolidation. This is also indicated indirectly by the recent results obtained in the clinic on patients with refractory epilepsy (Kragel et al., 2020); these results implicate increased theta/high gamma phase-amplitude coupling in the updating of previously formed memory traces.

It remains unclear whether similar or different mechanisms trigger theta-modulated slow and fast gamma rhythms. Obviously, more studies are needed to better understand the functional significance of gamma oscillations in the cortical-hippocampal network. Whatever the case may be, we believe that stronger theta/fast and theta/slow gamma coupling in the hippocampus and neocortical structures when rats explored of previously known objects indicates the retrieval of stored memory and encoding of new information.

### Successful object recognition depends on the difference in the strength of theta/gamma coupling during exploration

In our experiments, in most cases we observed that the strength of theta/gamma coupling in rats exploring the objects with novelty properties negatively correlated with the success of their discrimination. We propose that the successful recognition the object location and its temporal order depended on the *difference* in theta/gamma coupling during exploration of the objects with different properties *rather* than on its increase or decrease. The analysis of the correlation between the success in performing the *when/where* tasks and differences in the modulation index of gamma oscillations showed a pronounced correlation between theta/slow gamma and theta/fast gamma coupling in the hippocampus and the success in performing the *when* task. We also showed a correlation between the success in performing this task and the theta/slow gamma modulation in the neocortical structures. For the *where* component, theta/gamma coupling (for slow and fast gamma) correlated with ΔMI in the mPFC in a similar manner. In the hippocampus, this dependence was significant for the slow gamma band only. Interestingly, only in the mEC, the theta/gamma coupling in rats exploring of the displaced object was increased, and ΔMI negatively correlated with the success in the performance of the *where* task. This means that the higher phase–amplitude modulation in this structure reflects a higher probability of novelty recognition, but not the reactivation of previously acquired memory, as it was suggested for other structures. It has been hypothesized earlier that an increase in the theta/gamma coupling would serve to facilitate memory encoding in the entorhinal-hippocampal network (Colgin, 2016). Thus, the enhancement of theta/gamma coupling in the mEC may indicate the coding of new object placement in healthy rats.

In epileptogenic animals, the performance score in the *when* task correlated with ΔMI (A1-B1) to the same degree as it was in control animals. However, disturbances in the correlation of the theta/gamma coupling with the degree of discrimination of the object displacement compared to the control were observed. Namely, theta/slow gamma coupling in the hippocampus and neocortical areas did not reliably correlate with successful identification of the object displacement, though theta/fast gamma coupling in the hippocampus showed a correlation with the success in the performance of the *where* task.

There is evidence in the literature that different cortical inputs converge on the hippocampus to create unique contextual meanings (Morris, 2001; Davachi, 2006; Witter et al., 2000; Eichenbaum et al., 2007; Jo et al., 2007). Therefore, the role of the hippocampus in episodic memory should depend on its capacity to bind these attributes of unique experiences into a spatiotemporal representation. The theta rhythm is thought to promote the formation of long-term memories in the hippocampus by orchestrating inputs from cortical sources (MacDonald et al., 2011; Naya and Suzuki, 2011). Gamma rhythms, as has been shown by multiple findings, play important roles in memory retrieval and encoding of new information. As believed, slow gamma oscillations promote the retrieval of stored memories (Tort et al., 2009; Shirvalkar et al., 2010; Bieri et al., 2014; Takahashi et al., 2014; Igarashi et al., 2014; *but see* Kemere et al., 2013), while the fast gamma rhythm facilitates the encoding of current signals (Newman et al., 2013), in particular, the encoding of spatial information (Brun et al., 2002; Fyhn et al., 2004; Hafting et al., 2005; Brun et al., 2008; Colgin et al., 2009; Kemere et al., 2013; Cabral et al., 2014; Bieri et al., 2014; Takahashi et al., 2014; *but see* Ainge et al., 2007; Montgomery and Buzsaki, 2007; Yamamoto et al., 2014). Thus, these hypotheses regarding fast and slow gamma functions in memory processing require further testing.

## Conclusions

In the present work, we have demonstrated alterations in theta and gamma rhythms, violation of theta coherence and theta/gamma coupling in the hippocampus and neocortical structures in epileptogenic rats. We also found that these alterations are associated with the loss of ability to recognize objects during the episodic-like memory test. The data obtained showed the failure of the hippocampus and neocortical structures to correct the coding and retrieval of the current signals during epileptogenesis. Therefore, information exchange between the hippocampus and neocortex was disturbed when epileptogenesis progressed.

The temporal coordination of brain processes, as reflected by oscillatory patterning, synchronization, and cross-frequency coupling, is believed to be of importance, and we can expect that the disturbance of these dynamic processes would lead to specific disturbances of cognitive or executive functions. These “rhythmopathies” can reflect the network malfunctioning and assist in specifying the diagnosis of several diseases, such as temporal lobe epilepsy. Here, we demonstrated that the impairment of cognitive behavior in epileptogenic rats was associated with abnormalities of the rhythms and their discoordination in three connected brain structures, the hippocampus, the medial entorhinal cortex and the medial prefrontal cortex. We believe that these data can be used for early diagnosis of temporal lobe epilepsy.

## Conflict of interest statement

Nothing declared.

## Acknowledgments

The work was carried out within the framework of the State Assignment of Institute of Theoretical and Experimental Biophysics, Russian Academy of Sciences (No. 075-00381-21-00), and was supported by the Russian Science Foundation (project No. 20-65-46035).

The authors are grateful to Svetlana Viktorovna Sidorova for comments and help in preparing the manuscript.

## Notes

### Competing Interest Statement

The authors have declared no competing interest.

